# Stable QTL for malate levels in ripe fruit and their transferability across *Vitis* species

**DOI:** 10.1101/2020.12.14.422701

**Authors:** Noam Reshef, Avinash Karn, David C. Manns, Anna Katharine Mansfield, Lance Cadle-Davidson, Bruce Reisch, Gavin L. Sacks

## Abstract

Malate is a major contributor to the sourness of grape berries (*Vitis* spp.) and their products, such as wine. Excessive malate at maturity, commonly observed in wild *Vitis* grapes, is detrimental to grape and wine quality and complicates the introgression of valuable disease resistance and cold hardy genes through breeding. This study investigated an interspecific *Vitis* family that exhibited strong and stable variation in malate at ripeness for five years and tested the separate contribution of accumulation, degradation, and dilution to malate concentration in ripe fruit in the last year of study. Genotyping was performed using transferable rhAmpSeq haplotype markers, based on the *Vitis* collinear core genome. Three significant QTL for ripe fruit malate on chromosomes 1, 7, and 17, accounted for over two-fold and 6.9 g/L differences in ripe fruit malate, and explained 40.6% of the phenotypic variation. QTL on chromosomes 7 and 17 were stable in all and in three out of five years, respectively. Variation in pre-veraison malate was the major contributor to variation in ripe fruit malate (39%) and their associated QTL overlapped on chromosome 7, indicating a common genetic basis. However, use of transferable markers on a closely related *Vitis* family did not yield a common QTL across families. This suggests that diverse physiological mechanisms regulate the levels of this key metabolite in the *Vitis* genus, a conclusion supported by a review of over a dozen publications from the past decade, showing malate-associated genetic loci on all 19 chromosomes.

## Introduction

Malate is ubiquitous across biological systems, including plants, where it is a key intermediate in essential biochemical pathways^1^. Malate is one of the predominant organic acids that accumulates in fleshy fruits^2^ and is thus a major determinant of fruit sourness, titratable acidity, and pH^3,4^. The sensory and chemical effects of malate can extend to products of fleshy fruits, such as wines produced from cultivated wine grapes, and malate is reported to be inversely correlated with wine grape quality^5^.

In the European wine grape (*V*. *vinifera)*, malate typically exhibits biphasic behavior over the growing season^6^. Prior to the onset of ripening (veraison), malate is synthesized mainly from sugars^7^, and accumulates at concentrations reaching 40 mg/g fresh weight in the vacuoles of immature mesocarp cells^8^, or 20 g/L in the entire berry^9,10^. After veraison, malate decreases by over 80% on a concentration basis^8–10^ through its metabolism, via respiration and alternative pathways^7,11^, and by dilution, i.e. the increase in fruit volume.

Interestingly, malate concentration and its developmental pattern varies strongly among *Vitis* species. Unlike *vinifera*, wild *Vitis* species (e.g. *riparia* and *cinerea*) show a lack of or diminished dissimilation of malate during ripening^12^, and malate in ripe berries of these species often exceeds 20 g/L^12,13^. These wild species are of considerable interest to grape breeders due to their resistance to both biotic and abiotic stresses^14–16^, but excessive malate (and thus sourness) in wild *Vitis* and their progeny have so far limited their commercial use.

To expedite and improve grape breeding, considerable efforts have been made in recent years to develop appropriate markers to permit marker assisted selection (MAS)^17–20^ for desirable traits, and in the past decade over a dozen quantitative trait loci (QTL)-mapping studies have been performed in grape on acidity-related traits including malate (e.g. ^21–24^). However, no outstanding locus that is both stable (across years) and conserved (across studies and genetic backgrounds) has been reported. One potential complication with previous QTL studies on malate is that phenotyping has typically only focused on malate concentrations at ripeness. However, as previously mentioned, malate concentration at ripeness is the consequence of three distinct processes (accumulation, dissimilation, and dilution). To date, only three studies attempted to separately detect loci associated with accumulation and final concentration ^22,25,26^ and no QTL studies have accounted separately for the effects of degradation and dilution. Additionally, several of the QTL studies reported limited segregation for malate in ripe fruit as compared to the sizable range that exists across the *Vitis* genus. This may have limited loci stability, especially due to the sensitivity of malate to harvest date and the environment^6,27^. Finally, studies utilized distinct genetic marker systems including simple sequence repeat (SSR)^24,28,29^, single nucleotide polymorphism (SNP)^22,24^, and genotyping by sequencing (GBS)^25,30^, hampering the ability to compare between associated loci, and introducing bias due to polymorphism within primer or probe sites, limited polymorphism across species, or genomic structural variations among germplasm^18,20^.

This study reports on an interspecific mapping family that exhibited strong and stable variation in malate at ripeness for five years. In the final year of the study, the separate contribution of accumulation, degradation, and dilution to malate in ripe fruit was used to understand the physiological basis of the identified QTL. A novel system of genetic markers targeting the core genome and reported to be transferable across the *Vitis* genus^20^ was utilized on an additional family with close genetic relatedness, to assess their use as a tool to improve QTL consistency across families. Our results were integrated with the accumulated literature to report on the most stable and conserved loci associated with malate concentration in ripe grape berries.

## Materials and Methods

### Plant material

A complex interspecific F_1_ family of 346 genotypes was generated by crossing the cultivars ‘Horizon’ (‘Seyval blanc’ × ‘Schuyler’, with *V. vinifera*, *V. labrusca*, *V. aestivalis* and *V. rupestris* background) and breeding selection Illinois 547-1 (*V. rupestris* B38 × *V. cinerea* B9)^31,32^. The family was first established in 1988 and further enlarged in 1996^17^, and the vineyard maintained by Cornell University in Geneva, NY. This family segregates for flower sex^32^ and ca. 180 individuals have perfect flowers and therefore bear fruit. An additional interspecific mapping F_1_ family was generated by crossing ‘Horizon’ with *V. rupestris* B38 in 2008 resulting in 215 perfect-flowered vines, of which 118 produced fruit for this study. The *V. rupestris* B38 × ‘Horizon’ and ‘Horizon’ × Illinois 547-1 families share ‘Horizon’ and *V. rupestris* B38 in both ancestries. In addition, the families are maintained at the same farm.

### DNA extraction and rhAmpSeq genotyping

DNA from each vine in this study was extracted as described in Hyma, et al.^17^. rhAmpSeq sequencing and marker design were previously conducted using multiple species as described in Zou, et al.^20^ and the families in this study were genotyped using the 2000 rhAmpSeq marker panel. rhAmpSeq sequencing data for the mapping family was initially analyzed using the Perl script *analyze_amplicon.pl* (https://github.com/avinashkarn/analyze_amplicon/blob/master/analyze_amplicon.pl), and later re-analyzed with an upgraded pipeline optimized for rhAmpSeq data analysis (https://bitbucket.org/cornell_bioinformatics/amplicon). Finally, using a custom Perl script, *haplotype_to_VCF.pl* (https://github.com/avinashkarn/analyze_amplicon/blob/master/haplotype_to_VCF.pl), the four most frequent haplotype alleles for each marker in the *hapgeno* file were converted to a VCF file, where each haplotype allele of a marker was converted to a pseudo A, C, G or T allele. The raw converted VCF file was imported in TASSEL (Trait Analysis by association, Evolution and Linkage) 5.2.51 software and the genotypes were imputed using the LD-kNNi imputation plugin also known as LinkImpute (v1.1.4) using the default parameters (High LD Sites = 30, Number of nearest neighbors = 10, and max distance between site to find LD = 10,000,000).

### Genotype quality control

Multidimensional scaling (MDS) and Mendelian error detection were performed as quality control (QC) analyses to identify vines in both families with genotyping errors due to self-pollination, pollen contamination, or mislabeling. For MDS analysis, a genome-wide pairwise identity-by-state (IBS) distance matrix was first calculated in TASSEL software using 1 – IBS, followed by the MDS analysis. The first two principal coordinates (PCOs) from the MDS results were graphically depicted in the R statistical software using the “ggplot2*”* package^33^. Similarly, the Mendelian error detection analysis was performed using the *mendelian* plugin in BCFtools^34^, where a VCF and pedigree file from each family were used as inputs. Vines that failed these two QC analyses were removed from the genetic map construction and QTL analyses.

### Genetic map construction

After imputation and QC, pedigree information and VCF files of each individual vine were used as inputs in Lep-MAP3 v.0.2 (LM3) to construct the genetic maps. The following LM3 modules and steps were used to construct the genetic maps: (1) *ParentCall2* module of Lep-MAP3 was used to call parental genotypes; (2) *Filtering2* module was used to filter distorted markers based on a χ^2^ (chi-squared) test and monomorphism (non-segregating); 3) *SeparateChromosomes2* module was used to split the markers into linkage groups (LG) (4) *OrderMarkers2* module was used to order the markers within each LG using 20 iterations per group, and computing parental (sex specific) and sex averaged genetic distances. The marker order of the genetic maps was evaluated for the consistency, genome organization, and structural variation by correlating with their physical coordinates on the *V. vinifera* ‘PN40024’ reference genome (version 12X.v2). Finally, the phased output data from the *OrderMarkers2* step were converted into phased genotype data using the *map2genotypes.awk* script, which was further converted into 4way-cross format (where, “1 1” = 1, “1 2” = 2, “2 1” = 3, and “2 2” = 4) for the QTL analysis in the “R/qtl” package^35^ of R statistical software^36^.

### Sampling

The ‘Horizon’ × Illinois 547-1 family was sampled during five growing seasons, between 2011-2013 and 2018-2019, consisting of 93, 162, 142, 139, and 148 fruiting genotypes, respectively. Year to year variation in the number of fruiting genotypes was largely caused by bird damage, most prominently in 2011. In 2011-2013 and 2018, sampling was performed on the same date for all fruiting genotypes to represent the ripe fruit stage. The sampling date for each year was determined based on fruit color, in-field measurement of total soluble solids on a sub-sample of the progeny, and the sanitary state of the vineyard. The median total soluble solids content was 19.3, 21.3, 20.2, 17.7, and 19.9 °Brix, in 2011, 2012, 2013, 2018, and 2019, respectively.

In 2019, the ‘Horizon’ × Illinois 547-1 family was sampled at both pre-veraison (softening green berries) and post-veraison time points, and a modified sampling protocol was employed to better account for variation in phenology within the family. The date of onset of veraison was monitored across the family by manual and visual inspection of fruit softening and appearance of red color. Softening green berries were sampled from each fruiting genotype to represent the peak of fruit malate concentration. Additional sampling of the ripe fruit stage was performed 35-40 days following the first sampling date. This sampling approach allowed the study of malate accumulation and dissimilation in each genotype, and to assess the range of phonological variability across the family and its potential effect on the phenotypic data measured in preceding years.

The *V. rupestris* B38 × ‘Horizon’ family was sampled and analyzed as described for ‘Horizon’ × Illinois 547-1, in the 2012 and 2013 growing seasons. 118 and 116 genotypes were sampled in 2012 and 2013 and the median total soluble solids content at sampling was 22.8 and 18.6 °Brix, respectively.

All samples were collected in Ziploc bags, transported in a cooler box (approx. 10 °C) to the lab and stored at −80 °C until processing.

### Analysis of fruit composition

From each sampled genotype, 50 berries were weighed, crushed, and left for 10 min at room temperature to allow for skin tissue to macerate with the juice. The juice was filtered using a double layer of cheesecloth into 50 mL falcon tubes and centrifuged for five min at 5,000 RPM (Labnet Z400, Hermle Labortechnik, Wehingen, Germany). Total soluble solids were measured during 2011-2013 and 2018 using a digital refractometer (Sper Scientific, Ltd., model # 300053, Scottsdale, AZ). In 2011-2013, malate, tartrate, and citrate were quantified on an Agilent 1260 HPLC system (Santa Clara, CA) fitted with a Bio-Rad Aminex HPX-87H ion exclusion column (Hercules, CA) fitted with a micro-guard cation-H refill cartridge (Bio-Rad cat #125-0129) and UV/VIS diode array detector monitoring at 210 nm, based on a previously described method^37^. Standards (containing 1 g/L of each organic acid) and blanks were run every 10 analyses. In 2018 and 2019, glucose, fructose, and malate were quantified enzymatically (GF 8363 for sugars and ML 8343 for malic acid) on an RX Monaco autoanalyzer (Randox Laboratories Ltd.). Control samples were repeated every 30 to 40 samples and values had an average coefficient of variance of 3.7, 1.8, and 2.4% for glucose, total hexose sugars, and malate, respectively.

### Quantitative contributions of accumulation, dilution, and degradation to malate concentration during ripening

Data collected during 2019 from the ‘Horizon’ × Illinois 547-1 family included berry weight and malate concentration measured at the onset of veraison (i) and at ripeness, i.e. 35-40 days post-veraison (ii). Based on these data, an equation was constructed to separately express the contribution of malate accumulation, degradation, and berry dilution to malate concentrations measured at ripeness (ii), for each genotype. The concentration of malate at ripeness (ii) was expressed as the loss in malate concentration from the end of the accumulation stage, i.e. the onset of veraison (i), to ripeness (ii), caused by degradation and dilution as follows:

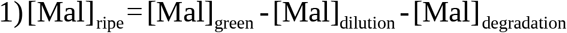

Where [Mal]_green_ and [Mal]_ripe_ represent the concentration of malate in the fruit (mg/mL) at sampling points (i) and (ii), respectively.

[Mal]_dilution_ represents the change in malate concentration between sampling points (i) and (ii), resulting only from the change in fruit volume:

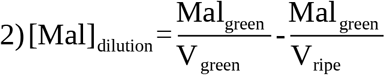

Where: V_green_ and V_ripe_ represent berry volume (mL) at sampling points (i) and (ii), respectively, and Mal_green_ represent malate content (mg/berry) at sampling point (i). Berry volume was calculated from the berry weight and specific gravity (S.G. 20/20), where S.G. was calculated from total soluble solid (TSS) content (°Brix) using standard sucrose conversion tables.

[Mal]_degradation_ represents the change in malate concentration between sampling points (i) and (ii) due to a change in malate content (in mg) per berry:

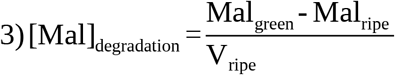

Where Mal_ripe_ represents malate content (mg/berry) at sampling point (ii).

### QTL analysis

Sex-averaged 4-way-cross genetic maps were used in QTL mapping using the “R/qtl” package^35^ of R statistical software^36^. The genotype probabilities were calculated using *calc.genoprob* with step = 0 (probabilities were calculated only at the marker locations) and assumed genotyping error rate of 1.0e-4. The *scanone* function and Haley-Knott (HK) method was used for QTL scan, and 1,000 permutation tests at alpha of 0.1 and 0.05 were performed to determine the significant LOD thresholds for each trait analyzed. Phenotypic data found to be non-linear or skewed were power transformed using *boxcoxnc* function in “AID” package. Further, QTL support intervals were determined calculating 1.5 – LOD support intervals using *lodint*. *Fitqtl* function was used to test for qtl-qtl interactions and calculate the total phenotypic variation explained by the QTL.

### Statistical analyses

The individual contribution of accumulation, degradation, and dilution to final malate concentrations was assessed using two statistical approaches: 1) Performing three Type I ANOVA tests (see equation 1), each with a different component, i.e. accumulation, degradation, and dilution, listed last. These tests are sequential, hence the effect of the last component listed is calculated after the effects of the previous two components have been controlled. ANOVA was run in R using basic *anova* and *lm* functions. 2) Performing a multilinear regression using equation 1 as a perfect model. For this, the package “hier.part”^38^, compatible with perfect models, was used. The relative contribution of each component was calculated by performing all regressions on the model components using *all.regs* function, followed by the function *partition*, which uses those regressions to partition the individual contribution of each component.

The broad-sense heritability (*H*^2^) was estimated by REML (restricted maximum likelihood) variance components method^39–41^ using linear mixed effects models implemented in the R package *lme4^42^* as 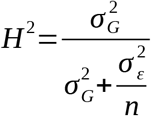 where *σ^2^_G_* and *σ^2^_ϵ_* are the variance of the genetic and residuals variance, respectively, and *n* is the number of experiments over the years, respectively.

Tukey HSD test was performed to compare the means of different haplotypes, using the built in *anova* function and *HSD.test* function from package “agricolae”^43^. Density plots were visualized using R package “ggplot2”^33^ and correlation matrices were based on Pearson’s correlation, performed using *chart.Correlation* in package “PerformanceAnalytics”^44^. MapChart^45^ was used to construct the physical map of chromosomes of interest.

## Results

### Progeny of two families covered a wide range of ripe fruit malate and were stable across multiple years

#### ‘Horizon’ × Illinois 547-1 family

Between 93 and 162 individual vines were characterized for fruit malate levels (g/L) at ripeness in the ‘Horizon’ × Illinois 547-1 family, during each of the five years of study (Fig. 1 A). Distribution for the data collected in 2011, 2013, and 2018 were skewed to the right. The family exhibited a mean five-fold difference in malate levels at ripeness each year, with values ranging between 3 and 21 g/L across years. These values are comparable to malate concentrations measured in *V. vinifera* cultivars^46^ and wild *V. cinerea* genotypes^12,13^, representing the low and high-end, respectively, both present in the family’s lineage. They cover approx. 70% of the entire range measured for malate in ripe fruit within the *Vitis* genus^13^.

**Figure 1.**
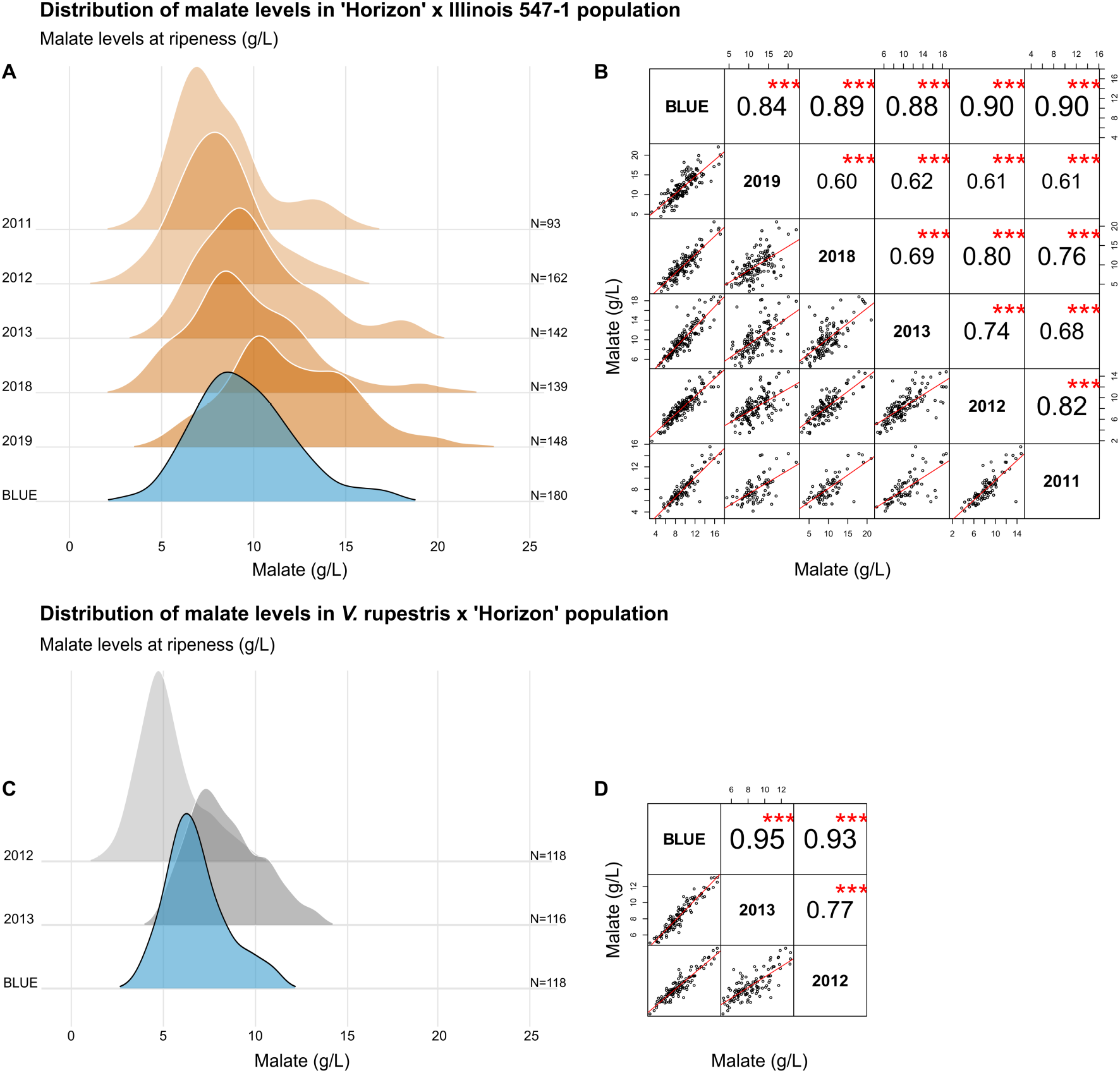
Malate concentrations (g/L) at ripeness in two interspecific *Vitis* families. The families were generated from a cross between ‘Horizon’ and either Illinois 547-1 or *V. rupestris* B38. Data was collected for five years (2011-2013, 2018-2019) for ‘Horizon’ × Illinois 547-1 and for two years for *V. rupestris B38* × ‘Horizon’ (2012 and 2013), and Best Linear Unbiased Estimates (BLUE) were calculated for each individual vine based on the multi-year data. **A, C)** Density plots representing the distribution of malate in each year and calculated multi-year BLUE for ‘Horizon’ × Illinois 547-1 and *V. rupestris* B38 × ‘Horizon’, respectively. **B, D)** Pairwise correlations for malate concentrations (g/L) measured in each year and the calculated multi-year BLUE. The upper-right diagonal panel shows pairwise Pearson’s correlation coefficients with their corresponding p values represented by asterisks, where ‘***’ stands for p<0.001. Bottom-left panel shows pairwise scatterplots, red lines represent best linear fit.

Differences in malate levels between progenies were stable from year to year and exhibited moderate-to-strong correlation between individual years (r=0.60 to 0.82), comparable to Chen, et al.^21^, and strong correlation with the calculated multi-year Best Linear Unbiased Estimates (BLUE, r=0.84 to 0.90) (Fig. 1 B). The broad sense heritability for malate concentration (g/L) at ripeness, calculated for five years of data, was 67%, similar to values reported by Bayo-Canha, et al.^24^ and slightly lower than those reported by Duchene, et al.^23^.

#### V. rupestris B38 × ‘Horizon’ family

The *V. rupestris* B38 × ‘Horizon’ family comprised 118 progenies, all characterized for fruit malate levels (g/L) at ripeness in 2012 and 2013, and a BLUE was calculated for the two years (Fig. 1 C). The data were also skewed to the right as in ‘Horizon’ × Illinois 547-1. Malate concentration at ripeness in the family ranged between 1.9 and 13.2 g/L in the two studied years, a narrower range of fruit malate compared to that measured for ‘Horizon’ × Illinois 547-1. Correlation was moderate to strong (r=0.77) between years and strong between years and the calculated BLUE (r=0.93 and 0.95) and indicated good year to year stability (Fig. 1 D). Broad sense heritability for malate concentrations at ripeness was 75%.

### rhAmpSeq genetic maps provide high-density mapping and transferability across *Vitis* spp

Out of 2,000 rhAmpSeq core genome markers 1,944 and 1,948 markers returned data for *V. rupestris* B38 × ‘Horizon’ (N=215) and ‘Horizon’ × Illinois 547-1 (N=346) families, respectively. Thirteen F1 vines from ‘Horizon’ × Illinois 547-1 and seven F1 vines from *V. rupestris* B38 × ‘Horizon’ failed QC analyses and were removed from the genetic map construction as contaminants (Supp. Files 1 and 2). Over 55% of the markers were placed on the genetic maps in 19 LG (Supp. Files 3 and 4), providing an average genome wide coverage of over 96%. The other 45% of the markers failed to be placed on the genetic map because they were either monomorphic or distorted. The total length of genetic maps ranged from 1050.7 to 1316.9 cM with an average marker density of 1.10 and 0.9 cM in *V. rupestris* B38 × ‘Horizon’ and ‘Horizon’ × Illinois 547-1, respectively. Similarly, marker orders on the genetic maps had a high correlation (r>0.94) to their physical coordinates (Fig. 2).

**Figure 2.**
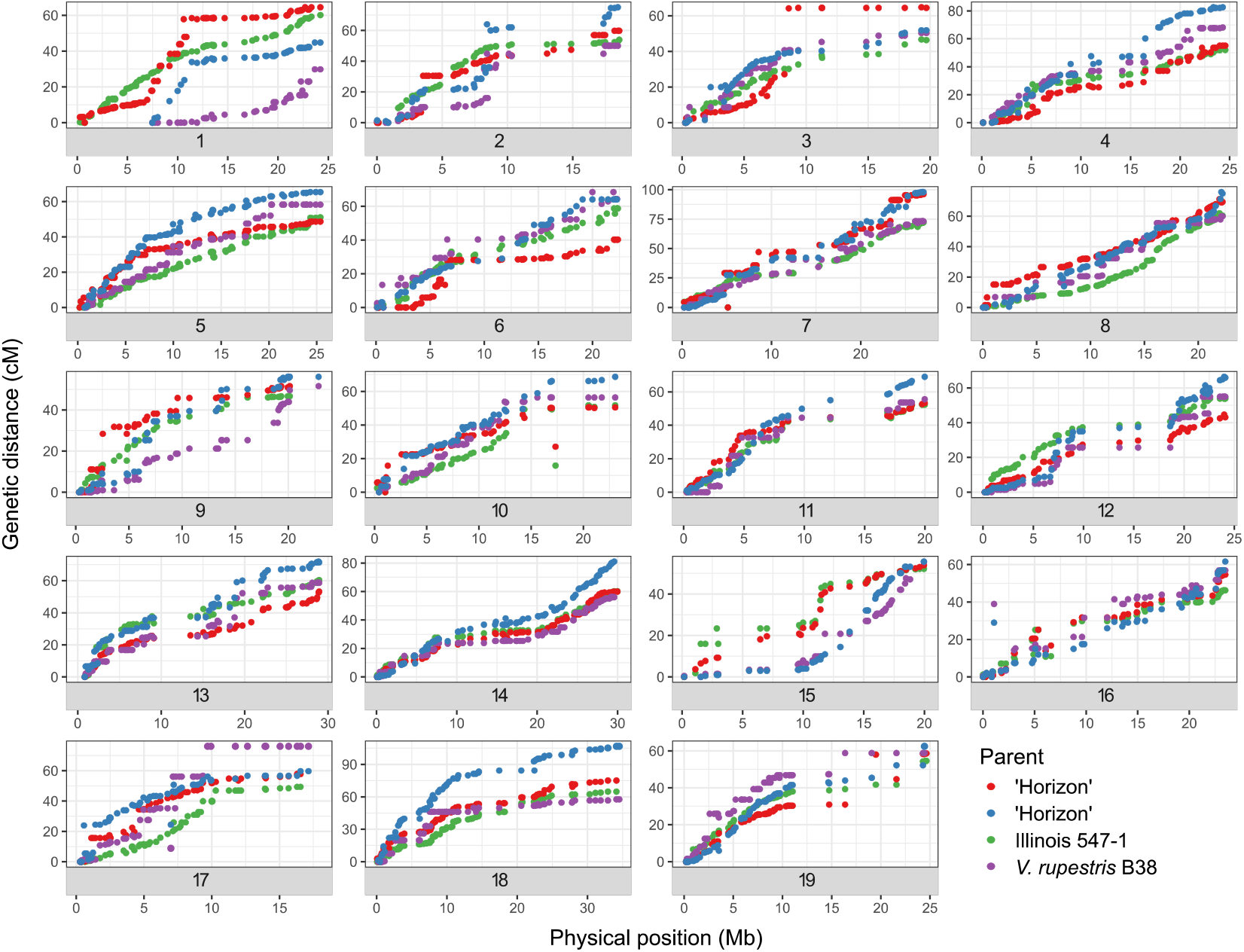
Relationship between the genetic and physical maps for the nineteen chromosomes of the parents of two interspecific grapevine families (‘Horizon’ × Illinois 547-1 and *V. rupestris* B38 × ‘Horizon’). Genetic distance: cM; Physical maps: Mb. The genetic distances of the markers were derived from the genetic map developed for each family and physical distances are from version 12X.2 of the PN40024 reference genome.

### Stable QTL on Chromosomes 7 and 17 for malate in ripe ‘Horizon’ × Illinois 547-1 fruit

QTL analyses of fruit malate concentrations at ripeness were performed separately on data from each of the five years sampled and from the best linear unbiased estimate (BLUE) calculated for these measured values. The data exhibited non-linearity and skewness in all years except for 2019 and was therefore transformed accordingly (Supp. File 5). A significant QTL on chromosome 7 was found in all five years of study and the BLUE (Table 1, Fig. 3). The position of the QTL shifted slightly among years but peaked at 15.37 Mb (44.2 cM) in 2012, 2013, and the BLUE (Table 1). An additional significant QTL on chromosome 17 was detected in 2012, 2013, 2018, and the BLUE, putatively overlapping with a malate QTL previously reported in a *vinifera* progeny^24^. The QTL on chromosome 7 explained a higher proportion of the phenotypic variance (mean=17.8%) than chromosome 17 (mean=12.6%). Two additional QTL were detected, on chromosome 10 in 2012 and on chromosome 1 in the BLUE but were not detected in other years. The significant QTL explained on average 31.3% of the total phenotypic variance measured in malate concentrations at ripeness across all years, and over 40% of the variance in 2012 and the BLUE, where three loci were significant. For BLUE values, both the combined effect of all three QTL (40.6%), and that of the two stable loci on chromosomes 7 and 17 (34.5%) constituted most of the broad-sense heritability of malate at ripeness (67%). No significant QTL-QTL interaction was detected in any of the years. The models were additive, and average malate levels at ripeness in the different haplotypes ranged from 5.5 to 12.4 g/L (n=5 and 6, respectively) in the calculated BLUE, accounting for more than a 2-fold difference in malate concentrations (Fig. 3 E).

**Table 1.** Summary of QTL associated with fruit malate concentration at ripeness in ‘Horizon’ × Illinois 547-1 and *V. rupestris* B38 × ‘Horizon’ families

| ‘Horizon’ × Illinois 547-1 | | | | | | | $H^2=0.67$ |
| --- | --- | --- | --- | --- | --- | --- | --- |
| Chr | Peak marker position (bp) | LOD Interval<sup>b</sup> (Mb) | Year | N | LOD score<sup>a</sup> | %variance QTL | %variance model<sup>c</sup> |
| 1 | 8,371,555 | 8.02-10.61 | BLUE | 164 | 4.44** | 6.12 | 40.6 |
| 7 | 10,066,816 | 7.5-23.4 | 2011 | 82 | 4.11* | 20.6 | 20.6 |
|  | 15,374,402 | 11.96-17.38 | 2012 | 147 | 7.64*** | 13.6 | 43.4 |
|  | 15,374,402 | 10.06-19.3 | 2013 | 130 | 5.77*** | 19.9 | 34.7 |
|  | 15,374,402 | 11.96 -20.3 | BLUE | 164 | 8.89*** | 17.8 | 40.6 |
|  | 18,952,564 | 11.96-20.3 | 2018 | 128 | 7.41*** | 21.8 | 35.5 |
|  | 22,781,295 | 11.96 -23.4 | 2019 | 137 | 4.2* | 13.2 | 13.2 |
| 10 | 17,335,086 | 2.6-17.35 | 2012 | 147 | 4.19* | 4.57 | 43.4 |
| 17 | 7,192,642 | 7.13-8.93 | 2012 | 147 | 5.89*** | 12 | 43.4 |
|  | 8,226,581 | 2.44-8.93 | 2013 | 130 | 4.54** | 16.3 | 34.7 |
|  | 8,226,581 | 7.29-9.35 | 2018 | 128 | 4.09* | 12.1 | 35.5 |
|  | 8,226,581 | 6.75 -8.93 | BLUE | 164 | 5.52*** | 10.1 | 40.6 |
| *V. rupestris* B38 × ‘Horizon’ | | | | | | | $H^2=0.75$ |
| 14 | 10,197,424 | 3.3-24.75 | 2013 | 111 | 4.28* | 9.03 | 24.98 |
|  | 10,197,424 | 3.3-26.07 | BLUE | 113 | 4.16* | 8.16 | 25.85 |
|  | 24,462,111 | 7.17-26.07 | 2012 | 113 | 4.30** | 10.6 | 26.9 |
| 15 | 17,991,582 | 16.81-19.96 | 2012 | 113 | 4.36** | 10.9 | 26.9 |
|  | 18,480,191 | 17.31-19.96 | 2013 | 111 | 4.19* | 10.89 | 24.98 |
|  | 18,480,191 | 17.22-19.96 | BLUE | 113 | 4.78** | 12.36 | 25.85 |

<sup>a</sup> LOD p values are represented by ‘\*’, ‘\*\*’, and ‘\*\*\*’ representing $p<0.1$ , $p<0.05$ , and $p<0.01$ , respectively, based on 1,000 permutation tests. <sup>b</sup> 1.5-LOD drop-off confidence interval in centimorgans (cM), <sup>c</sup> Additive QTL model combining all significant loci detected in the same year.

**Figure 3.**
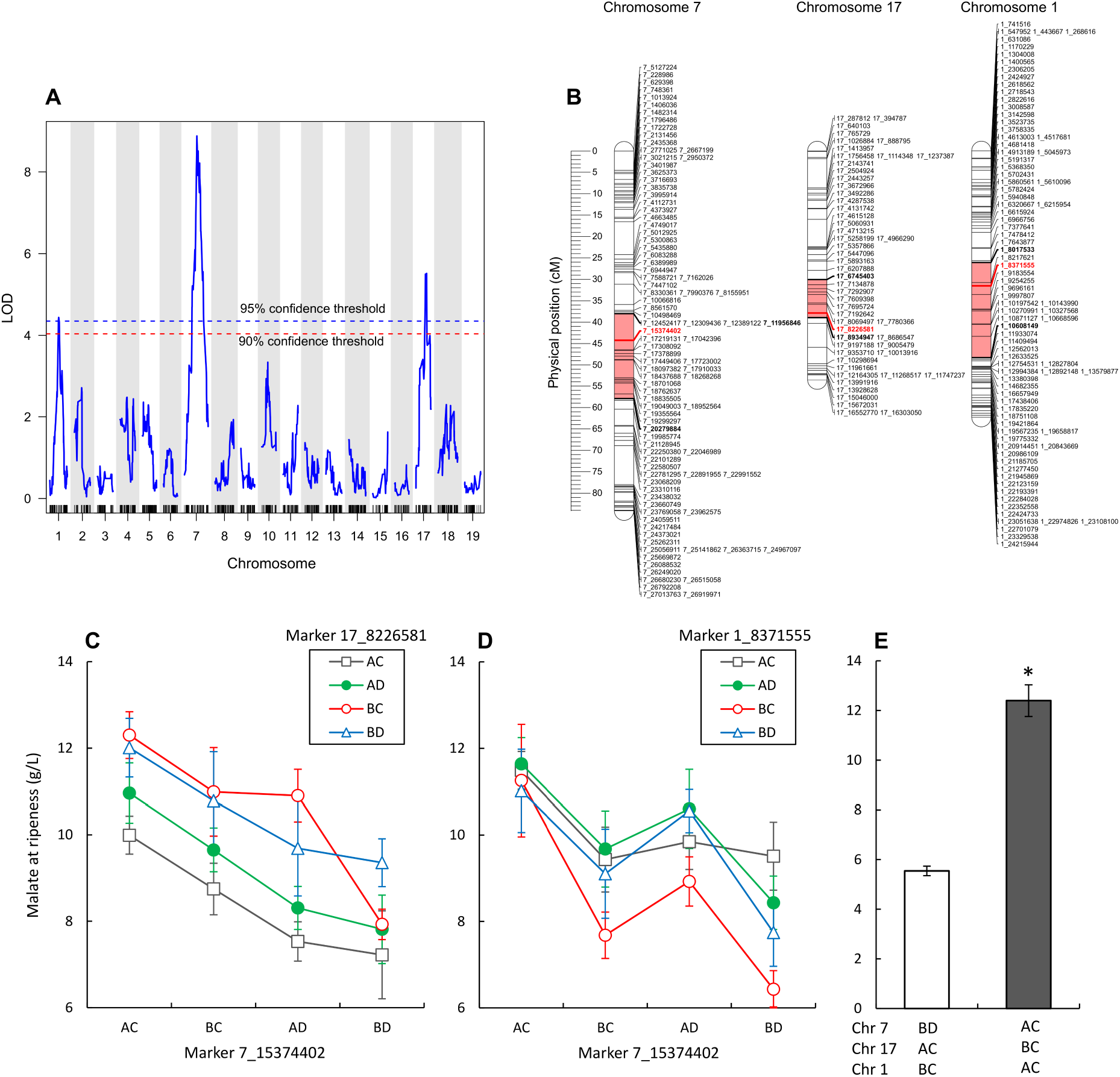
QTL for grape berry malate levels at ripeness. Data collected from an interspecific grapevine (*Vitis*) mapping family, generated from a cross between ‘Horizon’ and Illinois 547-1. QTL analysis was performed using the best linear unbiased estimate (BLUE) calculated on five-years of data (2011-2013, 2018, and 2019). **A)** Logarithms of the odds (LOD) score for genetic markers distributed across the 19 chromosomes. **B)** The physical position of peak markers (Bold red) and their LOD interval (interval is marked light red, edge markers are bold) on the three chromosomes found to significantly associate with the phenotype, ordered based on their relative contribution to the QTL model, from highest to lowest (7, 17, and 1, respectively). **C, D)** Effect of QTL marker haplotype combinations on fruit malate levels at ripeness, exhibiting the interaction between markers on chromosomes 7 and 17, and 7 and 1, in C and D, respectively. **E)** Haplotype combinations of all three loci yielding highest and lowest mean malate concentrations at ripeness (for n>2) in the family. n=6 and 5, respectively, asterisk represents statistically significant differences between means (p<0.01). Error bars are standard errors.

In the *V. rupestris* B38 × ‘Horizon’ family, all data were skewed and therefore transformed accordingly (Supp. File 5). Two stable QTL were identified in both years and the BLUE on chromosomes 14 and 15 and peaked at 10.2 and 18.48 Mb, respectively, in 2013 and the BLUE (Table 1). No significant QTL-QTL interaction was detected, and the additive model of these QTL explained up to 26.9% of the variance in ripe fruit malate. Average malate levels at ripeness in the different haplotypes ranged from 5.4 to 8.8 g/L (n=6 and 11, respectively) in the calculated BLUE (Supp. File. 6).

### Malate accumulation pre-veraison is the major factor determining ripe fruit levels

A set of equations separating the contribution of accumulation ([Mal]_green_), degradation ([Mal]_degradation_), and dilution ([Mal]_dilution_) on malate concentrations at ripeness ([Mal]_ripe_), were applied to the time-resolved data obtained in 2019 from the ‘Horizon’ × Illinois 547-1 family (materials and methods). Malate concentration at the onset of veraison was referred to as accumulation, as grape malate concentration reaches its peak at this stage prior to its typical dissimilation during ripening^10^. Dilution and degradation were calculated based on equation 2 and equation 3. Malate accumulation ranged from 12.4 to 31.7 g/L (Fig 4 A); Degradation and dilution ranged from (−2.5) to 17.7 and (−4) to 11.9 g/L, respectively. Negative dilution values may represent dehydration of the fruit at the final stages of ripening. Negative degradation values indicate that some proportion of the progeny continues to accumulate malate during ripening. This phenomenon was reported in accessions of *V. cinerea*, which is part of the lineage of the Illinois 547-1 parent^47^. In a pairwise correlation analysis (Fig 4 B), final malate concentration in the fruit (i.e. 35-40 days post-veraison) were positively correlated with malate accumulation (r=0.4) and negatively correlated with degradation and dilution (r= −0.21 and −0.33, respectively), as expected. In addition, degradation values were positively correlated with accumulation (r=0.56) and negatively correlated with dilution (r=-0.45). The former may indicate that high rates of malate degradation post-veraison are in part dependent on high pre-veraison malate concentrations.

**Figure 4.**
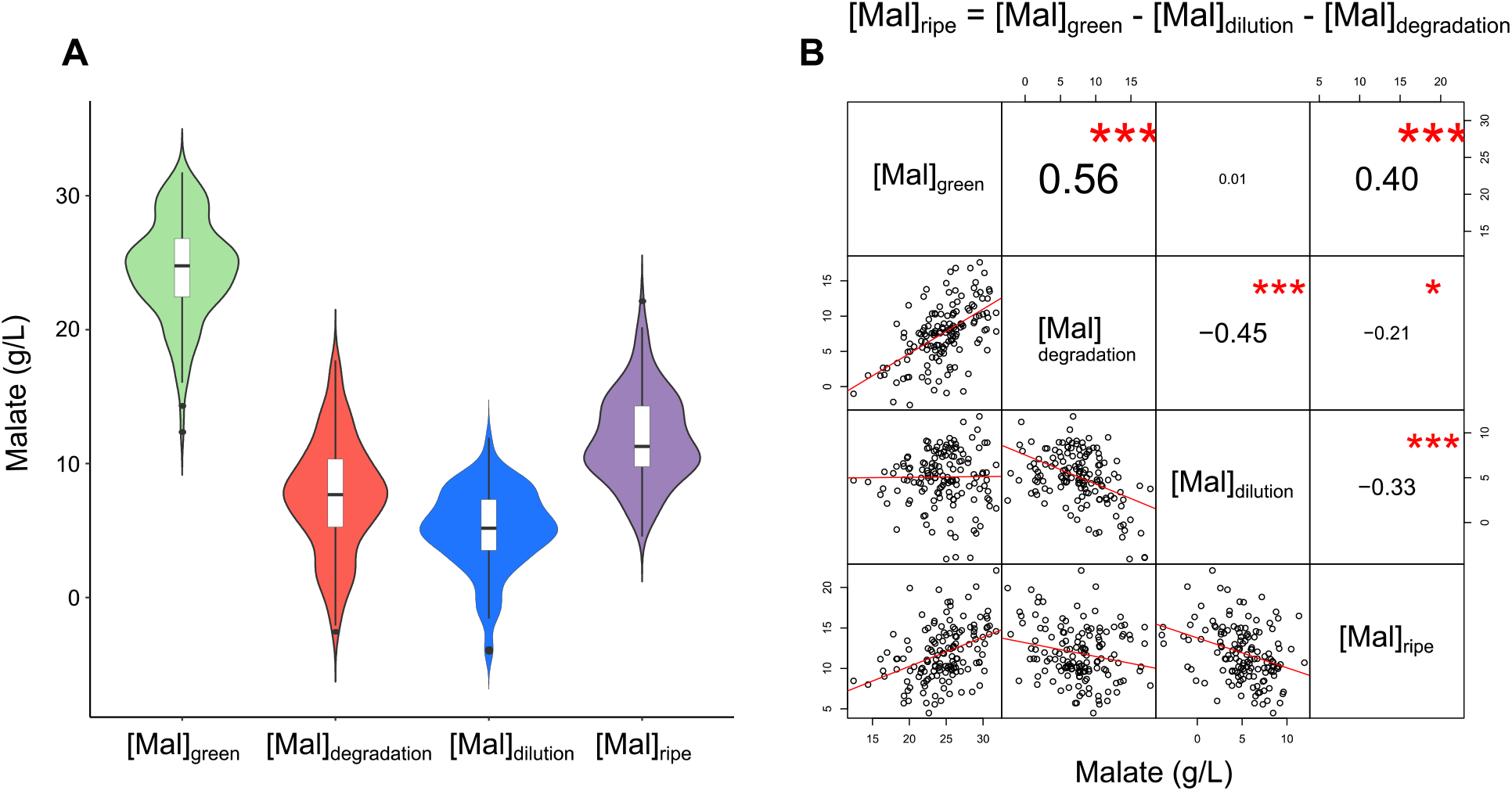
Measured and calculated components determining malate concentration in the fruit of an interspecific *Vitis* mapping family. The family was generated from a cross between ‘Horizon’ and Illinois 547-1. **A)** Malate concentrations at the onset of veraison ([Mal]_green_) and ripeness (35-40 days post-veraison, [Mal]_ripe_), measured in 2019, and the calculated concentration change due to degradation and dilution during the ripening stage ([Mal]_degradation_ and [Mal]_dilution_, respectively). **B)** Pairwise correlations of measured ([Mal]_green_ and [Mal]_ripe_) and calculated ([Mal]_degradation_ and [Mal]_dilution_) values. Upper-right diagonal panel shows pairwise Pearson’s correlation coefficients with their corresponding p values represented by asterisks, where ‘*’, ‘**’, and ‘***’ stand for p<0.05, p<0.01 and p<0.001, respectively. The bottom-left panel shows pairwise scatterplots, red lines represent best linear fit (n=148).

The individual contributions of accumulation, degradation, and dilution to the distribution of final malate concentrations in the family, were assessed using Type I ANOVA sum of squares and multilinear regression using equation 4 as a perfect model (Table 2). Both approaches indicated that accumulation, i.e. malate concentrations at the onset of veraison, explained the greatest proportion of the observed variation in malate at ripeness, but that the other two factors were also important contributors. Based on multilinear regression, accumulation explains 39% of the observed variation, followed by degradation (33.1%), and dilution (27.9%). The importance of all these factors to differences in grape malate at ripeness differs from observations on *Malus* (apple), where accumulation explains the majority of variation in malate at ripeness^47^.

**Table 2.**
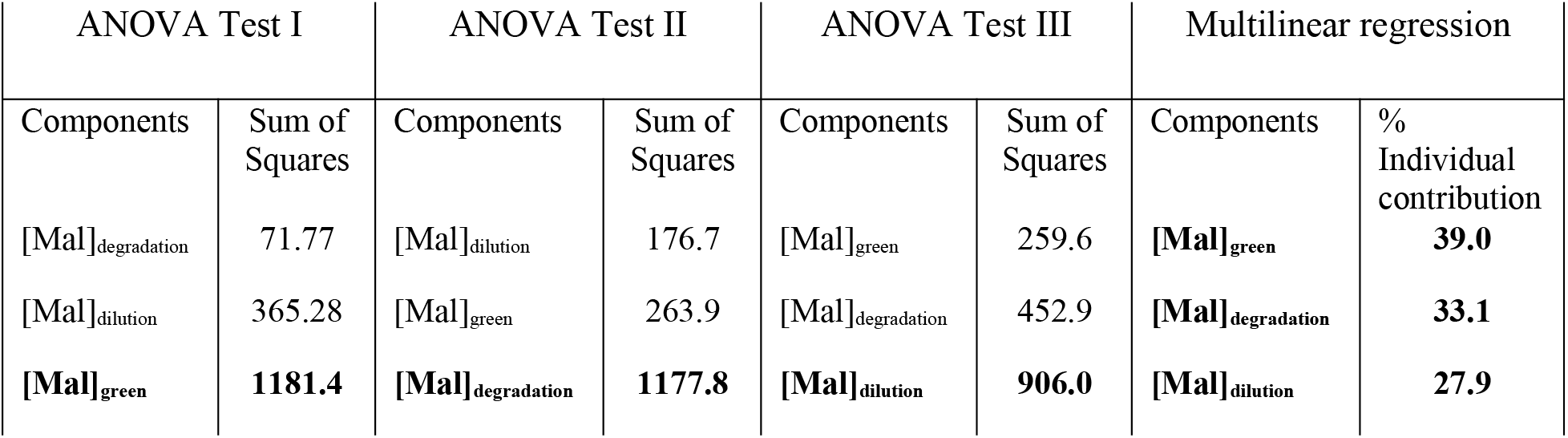
Type I ANOVA sum of squares and multilinear regression to assess the relative contribution of accumulation ([Mal]_green_), degradation ([Mal]_degradation_), and dilution ([Mal]_dilution_) on fruit malate levels at ripeness

### QTL for malate at pre-veraison and ripeness overlap on chromosome 7

Time-resolved data in 2019 facilitated the discovery of QTL associated with malate concentrations measured at the onset of veraison ([Mal]_green_) and ripeness ([Mal]_ripe_), and with malate degradation ([Mal]_degradation_), calculated as described in equation 3 (g/L). As reported above (Table 1), a single QTL on chromosome 7, peaking at 22.78 Mb (66.8 cM) was associated with malate concentrations at ripeness in 2019. Interestingly, an additional QTL on chromosome 7, peaking at 10.07 Mb (35.9 cM, LOD=5.18, p value=0.01) and explaining 16.4% of the variance, was found to be the only significant QTL associated with malate concentrations at the onset of veraison (Fig. 5 A, Supp. File 5). The LOD interval for malate accumulation overlaps with the QTL detected for ripe fruit malate in all years and the BLUE. This aligns with the important relative contribution of pre-veraison malate to ripe fruit malate in this family. A QTL on chromosome 7 for malate at the pre-veraison stage has been previously reported in grape^22^.

**Figure 5.**
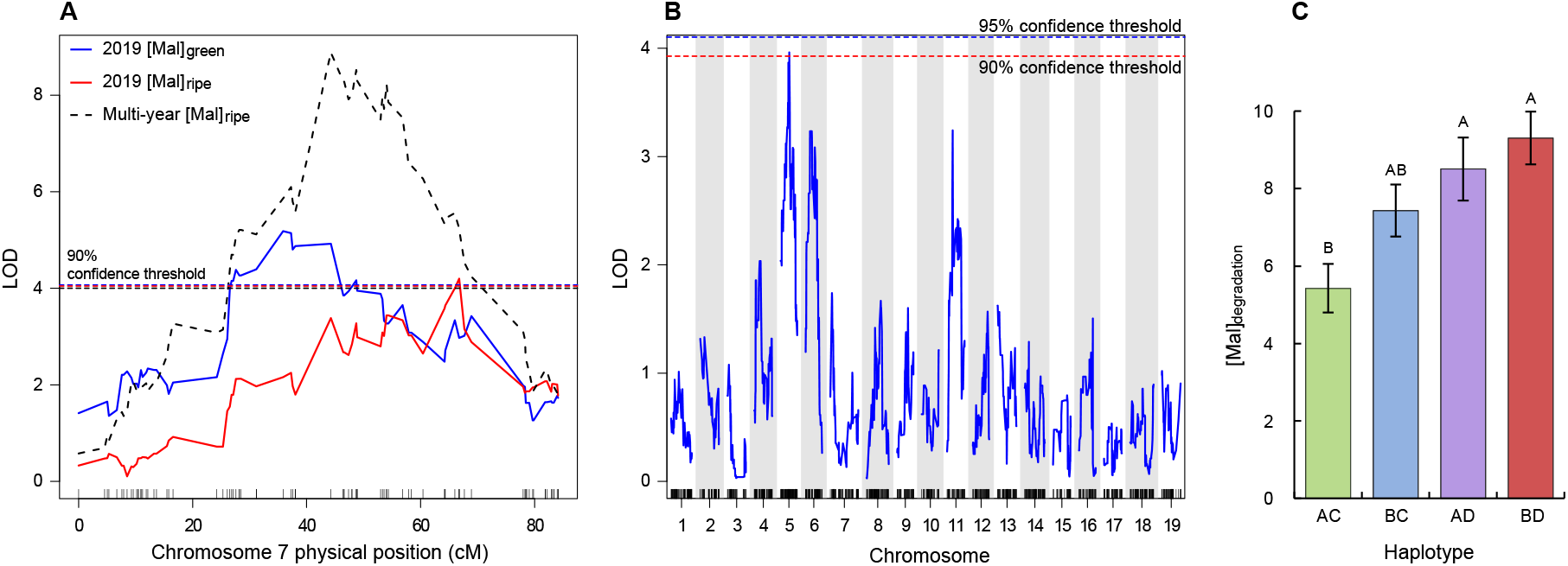
QTL analyses based on time-resolved malate concentration in the fruit, collected during 2019 from an interspecific *Vitis* mapping family, generated from a cross between ‘Horizon’ and Illinois 547-1. **A)** an overlay of the Logarithm of the Odds (LOD) score for malate concentrations (g/L) at ripeness and the onset of veraison in 2019 ([Mal]_ripe_ and [Mal]_green_, respectively), and the best linear unbiased estimate (BLUE) calculated for 5-year malate data at ripeness (Multi-year [Mal] _ripe_). Dashed horizontal lines represent the 90% confidence threshold for the phenotype marked with the corresponding color. N=148 for 2019 phenotypes, N=180 for multi-year BLUE. **B)** LOD scores for malate degradation ([Mal]_degradation_, g/L) during the ripening stage between the onset of veraison and ripeness (g/L). Red and blue dotted lines represent the 90 and 95% confidence threshold, respectively. N=148. **C)** Malate degradation ([Mal]_degradation_, g/L) calculated for the different haplotypes of the degradation QTL peak marker (5_9107670). Error bars represent the standard error. Bars marked by different letters represent significantly different values (Tukey test, p < 0.05). N=40, 35, 24, and 34 for AC, BC, AD, and BD, respectively.

A QTL on chromosome 5 (9.1Mb) (Fig. 5 B, C) was associated with the amount of malate degraded during ripening and explained 13% of the variance (Supp. File 5). The association was statistically weak, with a LOD score of 3.96 and p value of 0.097 based on 1,000 permutation tests. Average malate degradation during ripening in the different haplotypes ranged between 5.4 and 9.3 g/L accounting for approx. 4 g/L difference based on this QTL. Despite this marked effect, no QTL for malate at ripeness was found on chromosome 5 in any of the years of the current study, although a QTL on chromosome 5 for malate at ripeness has been reported for a *vinifera* family^24^ (Table 3).

**Table 3.** Summary of significant QTL associated with malate in ripe grape – current and previous works

| Chr/<br>LG | Years found<br>significant | Position (Mb or cM) | % Variance<br>explained | Authors |
| --- | --- | --- | --- | --- |
| **1** | **1/5**<br>2/2 | **8.37 Mb**<br>3.17-5.31 Mb | **6.1**<br>n.d. | **Current study**<br>Higginson <sup>25</sup> |
| 2 | 1/2<br>1/6<br>1/3 | 6.0/15.46 Mb<br>31.9 cM<br>n.d. | n.d.<br>10.1<br>10.1 | Marrano <sup>60</sup><br>Bayo-Canha, et al. <sup>24</sup><br>Schwander, et al. <sup>61</sup> |
| 3 | 1/3<br>1/3 | 10.94-12.21 Mb<br>n.d. | n.d.<br>9.8 | Higginson <sup>25</sup><br>Schwander, et al. <sup>61</sup> |
| 4 | 1/2<br>1/3<br>1/3 | 22.97 Mb<br>n.d.<br>69.6 cM | n.d.<br>12.1<br>11 | Marrano <sup>60</sup><br>Schwander, et al. <sup>61</sup><br><sup>b</sup> Houel, et al. <sup>22</sup> |
| 5 | 1/6 | 41 cM | 10.5 | Bayo-Canha, et al. <sup>24</sup> |
| 6 | 1/3<br>1/1 | 1.59 Mb<br>7.99 Mb | 17.31<br>26.19 | Chen, et al. <sup>21</sup><br>Yang, et al. <sup>30</sup> |
| **7** | **5/5**<br>2/6<br>1/3<br>1/3 | **15.37 Mb**<br>63.9/88.5 cM<br>n.d.<br>49/54 cM | **Up to 21.8**<br>Up to 11.9<br>8.4<br>Up to 15 | **Current study**<br>Bayo-Canha, et al. <sup>24</sup><br>Schwander, et al. <sup>61</sup><br><sup>b,c</sup> Houel, et al. <sup>22</sup> |
| 8 | 4/6<br>1/3 | 2.1-59.1 cM<br>15.23-15.32 Mb | Up to 20.4<br>n.d. | Bayo-Canha, et al. <sup>24</sup><br>Higginson <sup>25</sup> |
| 9 | 1/6 | 74 cM | 23.9 | Bayo-Canha, et al. <sup>24</sup> |
| **10** | **1/5** | **17.34 Mb** | **4.6** | **Current study** |
| 11 | 2/6<br>1/3 | 0/14.3-15 cM<br>n.d. | Up to 14.4<br>10.8 | Bayo-Canha, et al. <sup>24</sup><br>Schwander, et al. <sup>61</sup> |
| 12 | 1/2<br>2/3 | 3.45/7.13 Mb<br>n.d. | n.d.<br>Up to 14.7 | Marrano <sup>60</sup><br>Schwander, et al. <sup>61</sup> |
| 13 | 3/4 | 14-29 cM | up to 29.8 | <sup>a</sup> Ban, et al. <sup>28</sup> |
| **14** | **2/2**<br>1/2<br>3/3<br>1/6 | **10.2/24.54 Mb**<br>7.67 Mb<br>n.d.<br>13.5 cM | **Up to 10.64**<br>n.d.<br>Up to 25.5<br>14.7 | **Current study**<br>Marrano <sup>60</sup><br>Schwander, et al. <sup>61</sup><br>Bayo-Canha, et al. <sup>24</sup> |
| **15** | **2/2**<br>1/6 | **17.99/18.48 Mb**<br>56.5 cM | **Up to 12.36**<br>11 | **Current study**<br>Bayo-Canha, et al. <sup>24</sup> |
| 16 | 1/3 | n.d. | 13.9 | Schwander, et al. <sup>61</sup> |
| **17** | **3/5**<br>4/6 | **8.23 Mb**<br>0/26-30 cM | **Up to 16.3**<br>Up to 27.2 | **Current study**<br>Bayo-Canha, et al. <sup>24</sup> |
| 18 | 2/3 | 8.39/10.01/19.65 Mb | Up to 11.83 | Chen, et al. <sup>21</sup> |
| 19 | 1/2 | 6.33 Mb | n.d. | Marrano <sup>60</sup> |

QTL associated with phenotypes other than malate concentrations are marked by the following letters: a) titratable acidity, b) malate/total acids ratio, and c) malate/tartrate ratio. Position preferentially given in Mega base pairs (Mb) where available and alternatively in centimorgan (cM). n.d. – not determined. QTL detected by the current study are highlighted in bold text.

### Candidate gene identification

A total of 660 and 189 genes are found in the LOD intervals surrounding markers 7_15374402 and 17_8226581, respectively, stably associated with malate levels at ripeness. The LOD interval around marker 7_15374402 codes for several genes of interest involved in primary metabolism (phosphoenolpyruvate carboxykinase (PEPCK), acetyl-CoA carboxylase (ACC), NADH dehydrogenase, ATP synthase CF1 beta subunit), mitochondrial membrane trafficking (voltage-dependent anion channel 2 (VDAC2, marker position), dicarboxylate/tricarboxylate carrier (DTC)), phytohormones (small-auxin-up-RNAs (SAURs), auxin efflux carrier, Indole-3-acetic acid-amido synthetase (GH3.2), ethylene-responsive transcription factors (APETALA2/ERFs), ABA and brassinosteroid-regulators (AIP2, ABA 8’-hydroxylase, BRI1), and signaling and transcription regulation (protein phosphatase 2C (PP2C), protein kinases, and several MYB family transcription factors). Similarly, the LOD interval around marker 17_8226581 includes a cytosolic precursor of the mitochondrial malate dehydrogenase (mMDH), a major gene involved in malate metabolism and previously suggested as a candidate gene^24^. Additional candidate genes include a mitochondrial ATP synthase gamma chain, a mitochondrial carrier (exocyst subunit), several protein kinases (S6-kinase, part of Target of Rapamycin (TOR) signaling pathway), two MYB-related transcription factors, and a hypoxia-responsive gene.

Within the LOD interval for marker 7_10066816, associated with pre-veraison malate accumulation in 2019, we found 960 genes. These include the mitochondrial lipoamide dehydrogenase, a component shared among pyruvate dehydrogenase (PDH), 2-oxoglutarate dehydrogenase (OGDC), and branch-chained oxo-acid dehydrogenase (BCKDH) complexes, involved in the tricarboxylate cycle (TCA); two cytosolic malate dehydrogenase (cMDH), three NADH dehydrogenases, a vesicle-associated membrane transport protein (VAMP726); genes involved in hormonal signaling, including auxin-binding and auxin-sensitive (IAA12) proteins and ABA-insensitive 3, ethylene-responsive transcription factors, and cytokinin dehydrogenase, and several kinases including a group of Leucine-Rich Repeat receptor-like kinases (LRR-RLK). In addition, as the LOD interval of this QTL overlaps with the QTL for malate at ripeness (marker 7_15374402), they share all the candidate genes aforementioned for ripeness QTL on chromosome 7.

## Discussion

This five-year study of a grapevine family, expressing a broad range of malate concentrations, detected two stable loci associated with fruit malate concentration at ripeness. These loci, positioned on chromosomes 7 and 17, explained 40.6% of the phenotypic variation in this trait, and were detected in five, and three years of the study, respectively. Additional loci were detected for both ‘Horizon’ × Illinois 547-1 and the genetically related *V. rupestris* B38 × ‘Horizon’ families, and haplotype sequences associated with a low malate phenotype (expanding 1-2 Mb from identified QTL), were identified in both families (Supp. Files 7 and 8). Use of these sequences will facilitate MAS and the introgression of valuable traits from wild accessions to future cultivars, while controlling fruit malate levels at ripeness.

Owing to its typical biphasic behavior in grape, malate levels at ripeness represent the sum of temporally and physiologically distinct accumulation and dissimilation stages^6^. Therefore, in the final year of the study we performed time-resolved phenotyping at the onset of ripening and on ripe fruit to establish the relative contributions of accumulation, degradation, and berry expansion (dilution) on the variability of malate at ripeness. Differences among progenies in accumulation, degradation, and dilution reached 19.3, 20.2, and 15.9 g/L of malate, respectively, emphasizing the wide range of malate behavior across *Vitis* species, and specifically within the ‘Horizon’ × Illinois 547-1 family. Malate concentration at the onset of veraison was the major factor determining malate at ripeness, and the associated loci for both traits overlapped on chromosome 7. This indicates that the chromosome 7 locus detected in the multi-year study is likely involved in malate accumulation.

The synthesis of malate occurs mainly from sugars via the coupled pathways of glycolysis and the mitochondrial TCA cycle^7^. Accordingly, the QTL on chromosome 7 detected here and in a previous study^22^ codes for genes involved in mitochondrial activity, including major genes in the TCA cycle (PDH, OGDC), respiratory complex (NADH dehydrogenase, ATP synthase), and transport of energy and metabolites, including malate, across the mitochondrial membrane (VDAC2, DTC)^48,49^. Furthermore, two genes directly involved in malate synthesis and metabolism in the cytosol, cMDH and PEPCK, are found in the QTL.

Once synthesized, the transport of malate into the vacuole is energy dependent. The activity of tonoplast proton pumps maintains an electrochemical gradient that supports the facilitated diffusion of malate anion across the tonoplast^11^. Recent findings in citrus and apple highlight a complex regulatory network involved in this process. An ethylene-responsive factor (ERF) was found to interact with the vacuolar ATPase proton pump at the protein level, to modulate citric acid accumulation in citrus^50^. In apple, genotypic differences in malate accumulation were attributed to mutations in SAUR, PP2C, apple MYB transcription factor (MYB1), and aluminum-activated malate transporter (ALMT)^51,52^. These factors are suggested to interact and regulate the transport of malate across the tonoplast. Intriguingly, the grape pre-veraison malate QTL on chromosome 7 codes for all these regulatory components identified in apple, excluding ALMT, suggesting that their functional role in grape should be investigated. Moreover, the expression of PP2C, auxin-related genes (auxin-efflux carrier, GH3.2, IAA12), and two protein kinases, positioned in this QTL, is reported to be downregulated in grape post-veraison, at which point malate accumulation ceases^53^. Two other loci, recently reported to be associated with grape malate at veraison, were not detected in our current study. These QTL were located on chromosomes 6 and 8, which contain ALMT2 and vacuolar ATPase proton pump, respectively^26^.

Several genes located in the QTL detected for ripe fruit on chromosomes 7 and 17 are associated with the onset of ripening and with the dissimilation of malate. The expression level of APETALA2, positioned in the QTL on chromosome 7, was suggested as a key marker for the onset of grape ripening^54^. In *V. vinifera*, during that stage, malate is metabolized by several potential pathways, including gluconeogenesis^7^. This pathway is suggested to involve the transport of malate into the mitochondria by DTC^48^, the oxidation of malate to oxaloacetate (OAA) by mitochondrial malate dehydrogenase (mMDH), and the cytosolic conversion of OAA to phosphoenolpyruvate by PEPCK^55^. Accordingly, the expression pattern of these enzymes in *V. vinifera* grape agrees with their role in the dissimilation of malate^48,55–57^, and that of mMDH located in the QTL on chromosome 17, specifically, was found to be upregulated during ripening^57^. The co-localization of genes involved in malate biosynthesis and degradation, in a QTL that is associated with malate levels at the pre-veraison stage, may be regarded as confounding. However, as suggested by Sweetman, et al.^7^ and references therein, it is likely that both metabolic trends are active to some extent at every stage of berry development, and that changes in their rates determine the net change in malate during the pre- and post-veraison stages. In support of that, while PEPCK expression was upregulated at the dissimilation (i.e. post-veraison) stage, both its transcripts and enzyme activity were found to be present in malate-accumulating berries at the pre-veraison stage^7,55,58^.

### QTL for ripe fruit malate reported in Vitis spp. are scattered and unstable

The past decade has seen many investigations into the genetic regulation of acidity in grape, especially malate. Table 3 summarizes the findings of twelve studies conducted between the years 2013 and 2020, including the current one. Eight of these studies reported significant QTL for grape titratable acidity or malate at ripeness. These works included various genetic backgrounds, including a survey of *V. vinifera sylvestris* and *V. vinifera sativa* genotypes, pure *vinifera* families, and interspecific *Vitis* families. In summary, QTL associated with fruit malate or malate-dependent phenotypes (e.g. titratable acidity, malate/total acids ratio) have been reported in all 19 chromosomes of the *Vitis* genus. However, only 11 out of 37 QTL presented here (29.7%) were reported to be stable over at least two years and in at least 50% of the years tested. Other quality-related traits do not exhibit this lack of consistency. For instance, the presence of retrotransposons in VvMYB1 has been associated with non-pigmented skin color in multiple *Vitis* species^59^.

Potentially, some detected QTL could be artifacts resulting from an environmental effect, sampling bias, and other technical issues. However, considering that many of the reported QTL explain only a small amount of the variance in malate (<30%, and often much less; Table 3), it seems more likely that the instability of QTL across studies is a consequence of the large number of enzymes involved in the accumulation and dissimilation of malate^7^ and the likely involvement of a complex regulatory network^51,52^.

### Genetic markers are not a major source of inconsistency in malate QTL among families

Another possible explanation for the inconsistency of QTL across studies is that they vary in the technology and specific set of genetic markers used. We therefore tested whether using transferable rhAmpSeq markers would highlight common QTL associated with malate levels at ripeness in other interspecific families.

We observed significant QTL located at different chromosomes, even in families with strong genetic relatedness. The *V. rupestris* B38 × ‘Horizon’ and ‘Horizon’ × Illinois 547-1 families share ‘Horizon’ and *V. rupestris* B38 in both ancestries. The families were also grown at the same location (Geneva, NY). Nonetheless, significant QTL for multi-year values of malate at ripeness (BLUE) differed between the families and were located on chromosomes 14 and 15 for *V. rupestris* B38 × ‘Horizon’ compared to 7, 17, and 1 for ‘Horizon’ × Illinois 547-1 (Fig. 6). In addition, we tested for minor effects of allelic differences in the markers identified in one family on ripe fruit malate levels in the other. In both cases, these differences were not statistically significant (Supp. File 9).

**Figure 6.**
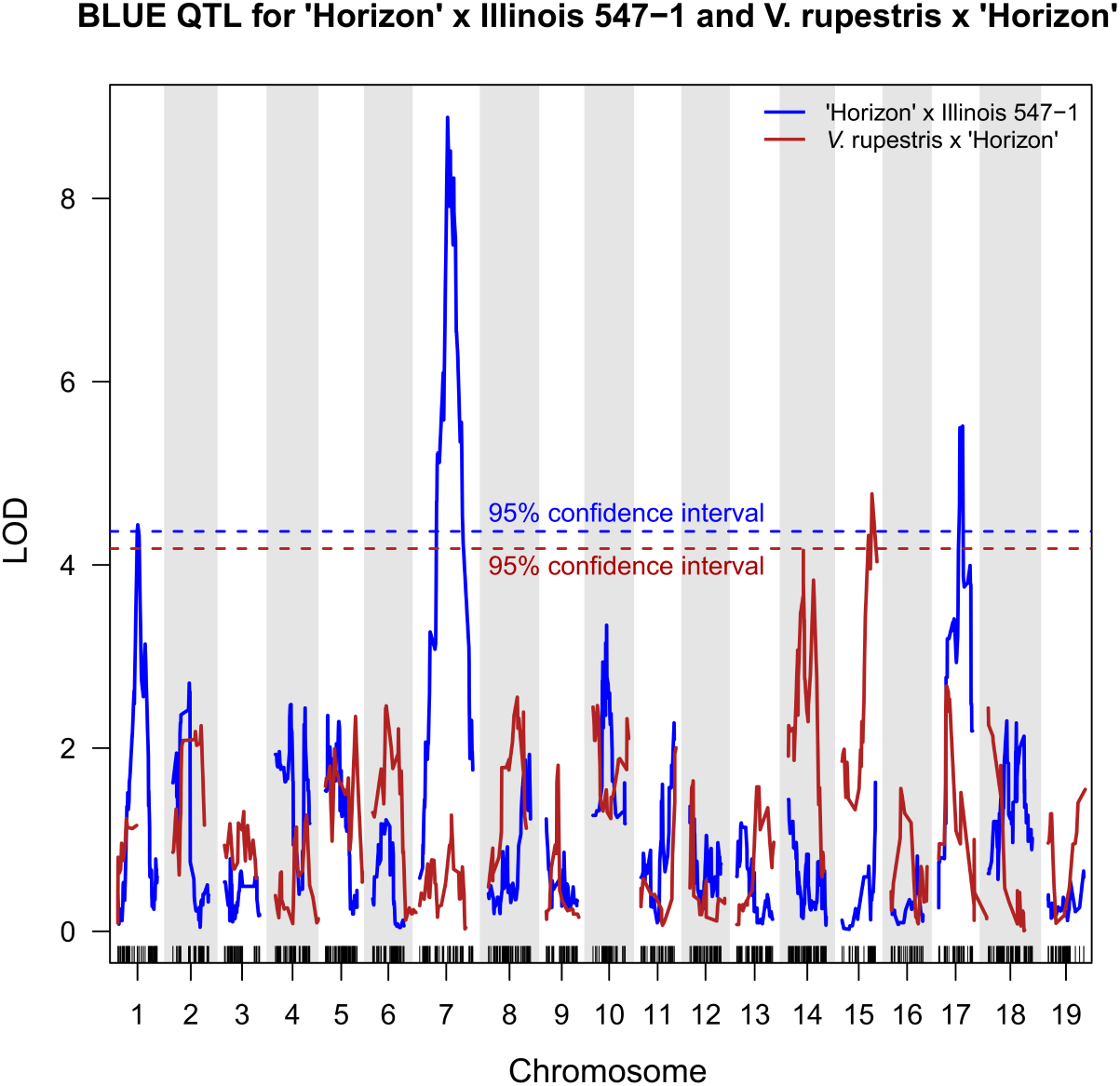
QTL for grape berry malate levels at ripeness in two genetically related interspecific grapevine (*Vitis*) mapping families. The families were generated from a cross between ‘Horizon’ and Illinois 547-1 and *V. rupestris* B38 and ‘Horizon’. QTL analyses were performed on the best linear unbiased estimate (BLUE) calculated on five and two years of data collected in these families, respectively, using the same set of transferable genetic markers. The figure shows the Logarithms of the odds (LOD) score for genetic markers distributed across the 19 chromosomes. Blue and red dashed horizontal lines represent the 95% confidence interval calculated for ‘Horizon’ × Illinois 547-1 and *V. rupestris* B38 × ‘Horizon’, respectively.

These results confirm that QTL for malate can be inconsistent even when using genetically related families studied in the same field and using the same, transferable, genetic markers^20^. This further suggests that the large variation in malate (~10-fold) in ripe fruit of the *Vitis* genus is a result of multiple enzymes directly involved in accumulation and dissimilation of malate and/or in a complex regulatory network. As such, the large number of QTL and inconsistency among previous studies likely represent different routes for control of malate levels in grape.

### Consistent and stable malate QTL across studies and diverse genetic backgrounds

To support the efforts for breeding cultivars with desired malate concentration, there is a need to identify loci that are stable from year to year, conserved between species, and have a strong impact on malate at ripeness.

Based on current literature, QTL located on chromosomes 1, 7, 8, 11, 12, 14, 15, and 17 were found stable, i.e. detected for several years in the same study, and highlighted by at least two independent studies. Of these, QTL on chromosomes 7 and 14 were highlighted by four independent studies, yet since the genetic locations of the markers have not been specified in most cases, it is not possible to determine if these loci indeed overlap. This lists the current putatively conserved QTL associated with grape malate at ripeness. Candidate genes suggested in these loci code for metabolic enzymes involved in the synthesis and degradation of malate.

Pyruvate kinase, on chromosome 8 (*VIT_08s0056g00190*)^24^, participates in the synthesis of malate from sugars through glycolysis^7^, isocitrate lyase and malate synthase on chromosomes 12 (*VIT_12s0059g02350*)^60^ and 17 (*VIT_17s0000g01820*)^24^, respectively, catalyze the synthesis of malate from isocitrate in the glyoxylate cycle, and mitochondrial malate dehydrogenase on chromosome 17 (*VIT_17s0000g06270*), suggested by Bayo-Canha, et al. ^24^ and the current study, catalyzes the interconversion of malate and OAA in the TCA cycle. Nonetheless, candidate gene identification is biased and involves the selection of genes with clear functional relevance out of a list of up to thousands of genes. Further functional research is required to improve our understanding of non-metabolic regulation of malate in grape and reveal additional candidate genes.

To distill major QTL of interest, it is essential to consider the impact of these QTL on fruit malate concentration. To the best of our knowledge, the markers detected here, offer the strongest combined effect on grape malate concentration at ripeness (6.9 g/L). As stable markers with high impact, they represent potential targets for further research and breeding efforts.

## Conclusion

This five-year study revealed two stable QTL associated with fruit malate levels at ripeness in an interspecific *Vitis* family. Combined, these QTL explain 40.6% of the phenotypic variation in the trait. Based on differences in malate levels between haplotypes in this family, utilization of the reported genetic markers in MAS is expected to accelerate the breeding of cultivars with improved palatability and provide source of variation for further fine mapping and molecular studies. Nonetheless, our analysis and the synthesis of existing literature highlights a lack of consistency and transferability of detected genetic markers, associated with malate, among *Vitis* families. This forms a major obstacle for the control of berry sourness through MAS. We hypothesize that the large number of molecular markers identified in this study and previous work, and their distribution across the grape genome, is a consequence of malate’s central role in primary metabolism. Therefore, control of grape malate and fruit sourness prompts better understanding of the intricate biological network and physiological processes involved in the regulation of malate. Studies of gene co-expression networks and alternative approaches to marker identification, such as eQTLs, may be more effective than conventional genomic QTL to provide the next breakthrough.

## Supplementary Files

**Supplementary files 1 and 2:** Genotyping QC for ‘Horizon’ × Illinois 547-1 and *V. rupestris* B38 × ‘Horizon’ families, respectively.

**Supplementary files 3 and 4:** 4waycross genetic maps for ‘Horizon’ × Illinois 547-1 and *V. rupestris* B38 × ‘Horizon’ families, respectively.

**Supplementary file 5:** Data transformation and QTL identified for malate model parameters.

**Supplementary file 6**: QTL effect plot and malate concentrations in haplotypes of *V. rupestris* B38 × ‘Horizon’ family.

**Supplementary files 7 and 8:** Haplotype sequences associated with low malate phenotype in ‘Horizon’ × Illinois 547-1 and *V. rupestris* B38 × ‘Horizon’ families, respectively.

**Supplementary file 9:** Effect of malate QTL markers found in ‘Horizon’ × Illinois 547-1 family on malate levels at ripeness in *V. rupestris* B38 × ‘Horizon’ family and vice versa.

## Funding

This research was funded by the US Department of Agriculture (USDA)-National Institute of Food and Agriculture (NIFA) Specialty Crop Research Initiative, award No. 2017-51181-26829.

## Acknowledgments

We thank the Cornell University Biotechnology Resource Center for providing the necessary equipment and technical support for the sequencing of DNA samples. We thank Mike Colizzi for the maintenance of the mapping families, and Heather Scott and Terry Bates for their support in the field.

## Competing interests

The authors declare that the research was conducted in the absence of any commercial or financial relationships that could be construed as a potential conflict of interest.

## Notes

### Competing Interest Statement

The authors have declared no competing interest.

