## Supplementary file 6 for "Stable QTL for malate levels in ripe fruit and their transferability across *Vitis* species"

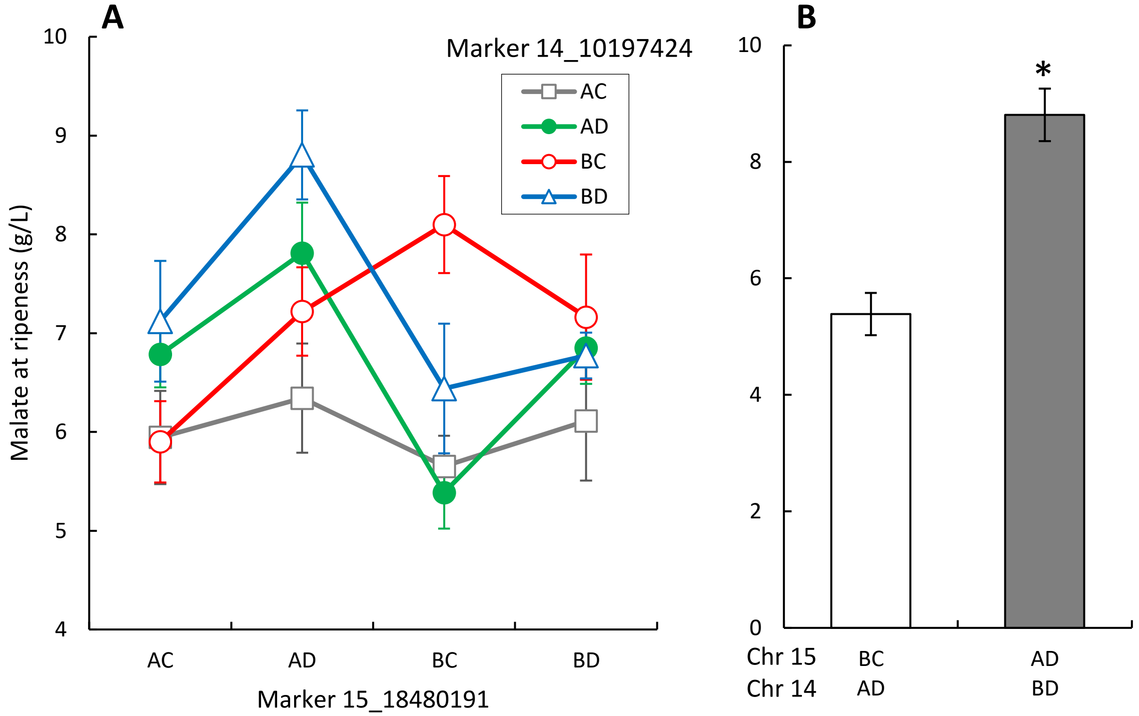


**Supplementary figure 1. QTL for grape berry malate levels at ripeness** in an interspecific grapevine (*Vitis*) mapping family, generated from a cross between *V. rupestris* and 'Horizon'. QTL analysis was performed using the best linear unbiased estimate (BLUE) calculated on two years of data (2012-2013). **A)** Effect of QTL marker haplotype combinations on fruit malate levels at ripeness, exhibiting the interaction between markers on chromosomes 14 and 15. **B)** Haplotype combinations of the two loci yielding highest and lowest mean malate concentrations at ripeness in the family, n=11 and 6, respectively. Error bars are standard errors. *Means significantly differ based on a Tukey HSD test, p value < 0.001.
