## Supplementary file 9 for "Stable QTL for malate levels in ripe fruit and their transferability across *Vitis* species"

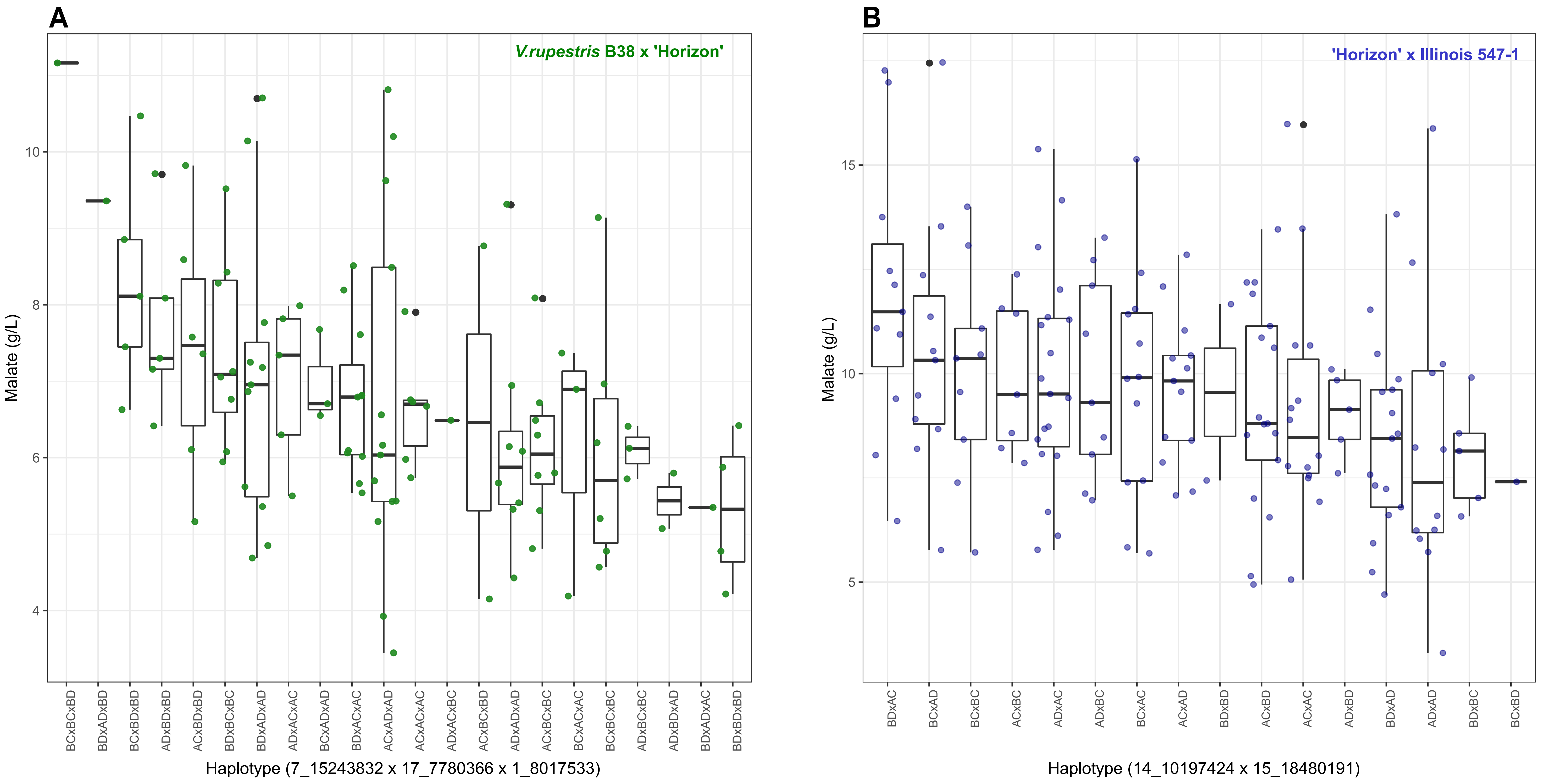


**Supplementary figure 2. Effect of malate QTL marker haplotypes found in one interspecific *Vitis* family on ripe fruit malate levels in a genetically related family. A)** Effect of haplotypes in QTL markers identified in 'Horizon' × Illinois 547-1 family on ripe fruit malate levels in *V. rupestris* B38 × 'Horizon' family. **(B)** Effect of haplotypes in QTL markers identified in *V. rupestris* B38 × 'Horizon' family on ripe fruit malate levels in 'Horizon' × Illinois 547-1 family. No statistically significant differences were found between groups based on ANOVA and Tukey HSD tests.
